# Non-linear tradeoffs allow the cooperation game to evolve from Prisoner’s Dilemma to Snow Drift

**DOI:** 10.1101/091041

**Authors:** Lin Chao, Santiago F. Elena

**Affiliations:** Section of Ecology, Behavior and Evolution, Division of Biological Sciences, University of California San Diego, La Jolla, California 92093-0116, USA; Instituto de Biología Molecular y Celular de Plantas (IBMCP), Consejo Superior de Investigaciones Científicas-Universidad Politécnica de Valencia, Ingeniero Fausto Elio s/n, 46022 Valencia, Spain; Instituto de Biología Integrativa y de Sistemas (I^2^SysBio), Consejo Superior de Investigaciones Científicas-Universitat de Valencia, Catedrático Agustín Escardino 9, 46182 Paterna, Valencia, Spain; The Santa Fe Institute, 1399 Hyde Park Road, Santa Fe NM 87501, USA

**Keywords:** Cooperation, Defective interfering particles, Game theory, Prisoner’s Dilemma, Snow Drift, RNA viruses

## Abstract

The existence of cooperation, or the production of public goods, is an evolutionary problem. Cooperation is not favored because the Prisoner’s Dilemma (PD) game drives cooperators to extinction. We have re-analyzed this problem by using RNA viruses to motivate a model for the evolution of cooperation. Gene products are the public goods and group size is the number of virions co-infecting the same host cell. Our results show that if the tradeoff between replication and production of gene products is linear, PD is observed. However, if the tradeoff is nonlinear, the viruses evolve into separate lineages of ultra-defectors and ultra-cooperators as group size is increased. The nonlinearity was justified by the existence of real viral ultra-defectors, known as defective interfering (DI) particles, which gain a nonlinear advantage by being smaller. The evolution of ultra-defectors and ultra-cooperators creates the Snow Drift game, which promotes high-level production of public goods.

## 1. Introduction

RNA viruses are ideal organisms for modeling the evolution of cooperation. Besides providing some of the more accurate information on the dynamics of the process, the molecular mechanisms that drive it have been elucidated to great detail. They are as simple as possible, but not simpler of a general case. All viruses are parasites that need to infect a host cell to reproduce. If a single virion infects a cell, kin selection is the agent of evolution because the progeny viruses are clonal. Kin selection may be uniquely important, if not actively enforced, in many RNA viruses because the first one or few viruses to infect a cell may exclude the entry of others, a phenomenon known as superinfection exclusion [1-3]. However, if a larger number of viruses are able to co-infect the same cell, groups of genetically unrelated viruses are created and opportunities for the evolution of cooperation and defection arise. The interaction between unrelated viruses in a mixed co-infection group creates payoff matrices at the fitness level that correspond to many standard game theory outcomes such as Prisoner’s dilemma (PD) and Snow Drift (SD; also known as the Hawk-Dove or the Chicken games) [4-9]. By increasing the group size of viruses co-infecting the same host cell, it has been possible to experimentally demonstrate that RNA viruses can evolve to be trapped by PD [10], yet escaping from this game is still possible if kin selection is restored [11].

A particularly appealing example of the interaction between cooperators and defectors in the Virosphere are replicator-defective mutants known as defective interfering (DI) particles and the wildtype viruses from which they evolve and depend for their replication and transmission [12-14]. Almost all viruses produce DIs, deleted forms of the genome of the wildtype virus, during replication. DIs are encapsidated into virus particles produced by a wildtype coinfecting virus and can be transmitted in a manner identical to the wildtype virus. Obviously, since DIs need the assistance of the wildtype virus to replicate and encapsidate, they can only persist in the long term at high multiplicity of infections, when more than one particle enters into susceptible cells [14,15]. Within coinfected cells, DI and wildtype virus genomes compete for resources, including binding to the viral replicase and packaging proteins. This competition results in a reduced accumulation of the wildtype virus, a process known as interference [13,16-25]. The interest on DIs has been revitalized in recent years mainly for three reasons: (*i*) they may be involved in triggering antiviral immunity during acute viral replication [26], (*ii*) they negatively impact the biotechnological production of vaccines and viral vectors [27] and (*iii*) their possible application as transmissible antivirals to control viral infections at the host-population level [28], the so-called therapeutic interfering particles.

One of the best studied DI-virus systems was the molecular control of replication and transcription in *Vesicular stomatitis virus* (VSV; genus *Rhabdovirus*, family Rhabdoviridae) (figure 1A), which we have taken as our idealized virus to model. This system offers a near perfect mechanism for modeling the tradeoff between defection and producing public goods [13]. The molecular control in VSV DI particles was also key for evaluating the non-linearity of the tradeoff in ultra-defectors (figure 1B).

**Figure 1.**
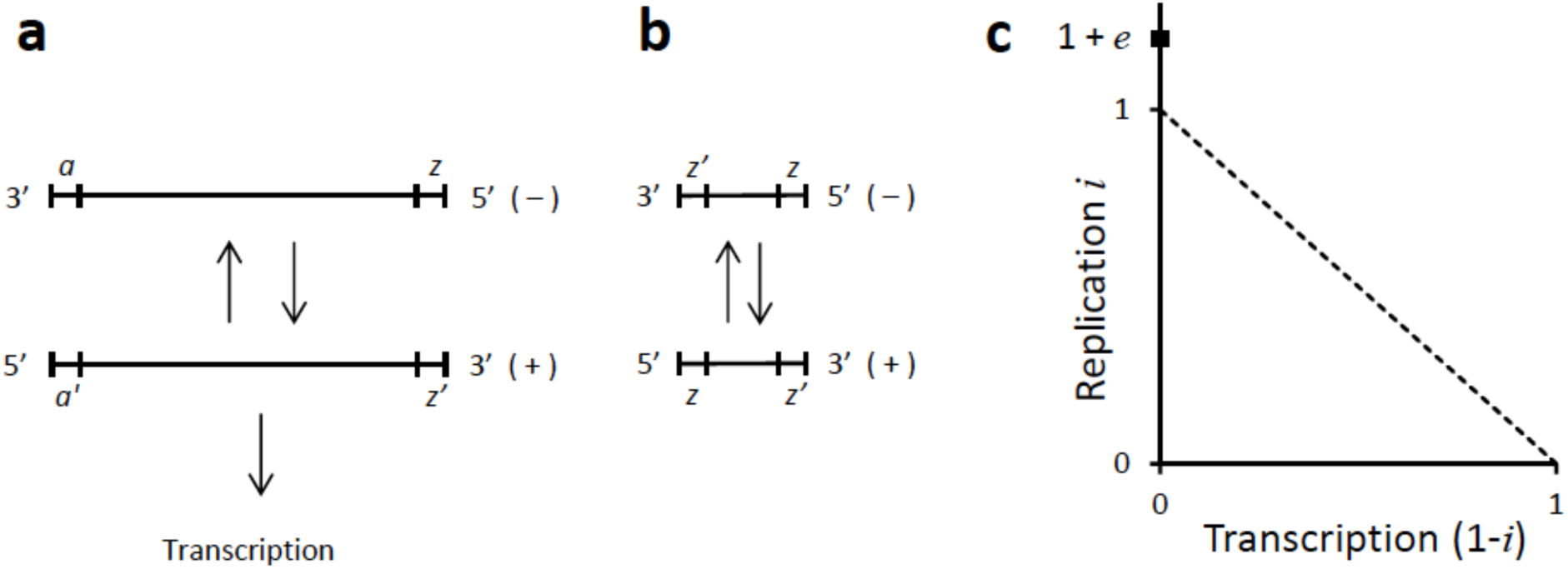
. Tradeoff between replication and transcription in VSV viruses and DIs. (**A**) Complete single stranded RNA genome of VSV with all required genes. After the (−) strand enters a host cell, the segment *z* serves as the initiation site for the synthesis of the (+) strand, which acts as both the messenger RNA and the replication template for the (−) strand. The segment *a’* acts as the initiation site for both transcription and replication and the (+) strand is therefore constrained to tradeoff between providing public goods and reproduction. (**B**) Single stranded genomes of DI particles. This shortened DI genome is the most abundant type and it lacks the coding regions for genes needed for replication and infection. Additionally, the (+) and (−) strands become functionally equivalent and only capable of replication because their *a’* and *a* segments are replaced with *z* and *z’* segments, respectively. (**C**) Linear and non-linear tradeoffs between replication and transcription in VSV. Following the model, a virus can allocate an amount of available resources *i* to replication and 1 − *i* to transcription. In the absence of DIs, a linear tradeoff (‐‐‐‐) is assumed between *i* and 1 − *i* because a virus can only tradeoff by modulating the initiation site *a’* to favor either replication or transcription, 0 ≤ *i* ≤ 1. If *i* = 1 and 1 − *i* = 0, *a’* has been modulated to promote only replication. With the evolution of DIs the tradeoff becomes nonlinear because DIs acquire an even higher replication by both foregoing transcription and being smaller and replication rate *i* > 1. To prevent *i* from becoming infinity large, a replication cap of 1 + *e* was set (■), where *i* ≥ 0 and *e* = 0 reverts to a linear tradeoff. Because *i* > 1 makes the transcription rate 1 − *i* negative, the tradeoff was bounded 1 − *i* ≥ 0. DIs with *i* = 1 + *e* > 1 were termed ultra-defectors.

## 2. Results and Discussion

Our model accommodates a viral population of size *N* that is randomly divided into *N/m* groups of size *m*≥1,where each group represents viruses that co-infect the same host cell. For virologists, *m* would correspond to multiplicity of infection (MOI) [29-31]. Once inside a cell, a virus must tradeoff between replicating its own genome and producing gene products. This aspect of the tradeoff was assumed to be linear because replication and transcription in VSV compete for the same initiation site (figure 1A). Thus, let *i* and 1 − *i* represent the effort an individual virus allocates to replicating its own genome and making public goods (*e.g*., proteins), respectively (with *i* ∈ [0,1]). We assumed additionally that an individual virus *n* in a group of size *m* has access to 1/*m* of the host’s resources and therefore is able to make only *i_n_/m* public goods and (1 − *i_n_*)*/m* genomes, where the subscript *n* is here and hereafter used to denote the effort by of the *n*^th^ virus and *n* = 1, 2, 3,…,*m*. This assumption is justified because many viral DIs, as per their name, interfere by reducing the total yield of wildtype and defective genomes produced by a co-infection group [16-25]. The reduction has also been shown to be linear, *e.g*. it is *i* = ½ when co-infections groups are 50% wildtype and 50% DI viruses [17]. After genomes and gene products are assembled into virions, a progeny of *b* number of viruses is released and the model determines the final fitness of an individual virus as

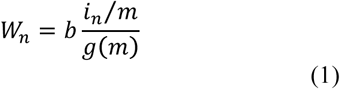
where 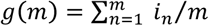 is the total number of genomes produced by the group of size *m*. To determine the value of b, we note that while its value could be equal to the total amount of public goods produced by the group, namely *h*(*m*)=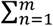(1-*i_n_*)*m* it could be less than *h*(*m*) if insufficient genomes are made. If both *g*(*m*) and *h*(*m*) are scaled in molar units required to assemble a viable virus, then *b* = min[*g*(*m*), *h*(*m*)]. If *g*(*m*) > *h*(*m*), the excess genomes are wasted and *h*(*m*) caps the number of viruses produced. *Vice versa* if *g*(*m*) < *h*(*m*) and then genomes are the limiting factor and public goods are produced in excess. Equation (1) is formally equivalent to the fitness equations developed by Frank [7,8] to study the evolution of parasites and protocells.

We modeled evolution by Monte Carlo simulations (see section 4a below) of a parent population of size *N* with known or assigned values of *i* and assembling *N/m* random groups. By using equation (1), the fitness of each virus was determined. A progeny population was then created by sampling *N* viruses from the parent population. Selection was imposed because the fitness values were used to weight the sampling process. The progeny was then re-assembled into *N/m* groups to create the parent population for the next generation. By using equation (1) and the weighted sampling, the process can be repeated to create new progeny and parent populations for as many generations as needed. If desired, the values of *i* were mutated before selection, in which case changes in the values of *i* over time in a population could be monitored to track the evolutionary dynamics of cooperation and to search for steady state outcomes.

### (a) Evolution is trapped by PD with a linear tradeoff

Used as presented above, equation (1) represents a linear tradeoff between replication *i* and transcription 1 – *i*. In a later section, non-linear tradeoffs are introduced by changing relationship of replication and transcription. A first test was to model evolution with *m* =1, for which an evolved optimum *i_k_* = ½ was anticipated, and observed, because each group is clonal and kin selection between groups favors viruses that make equi-molar amounts of genomes and gene products (figure 2A). Thus, we used *i_k_* = ½ as the evolutionary starting point and examined the effect of increasing *m*. In all cases *i* evolved dynamically to higher steady state values as *m* was increased (figure 2B). Because all of the populations remained monomorphic, we were able to determine analytically the steady state values by solving for the *i* that maximized individual fitness in a group of *mi* identical viruses (see section 4b below for the analytical derivation). The solution

**Figure 2.**
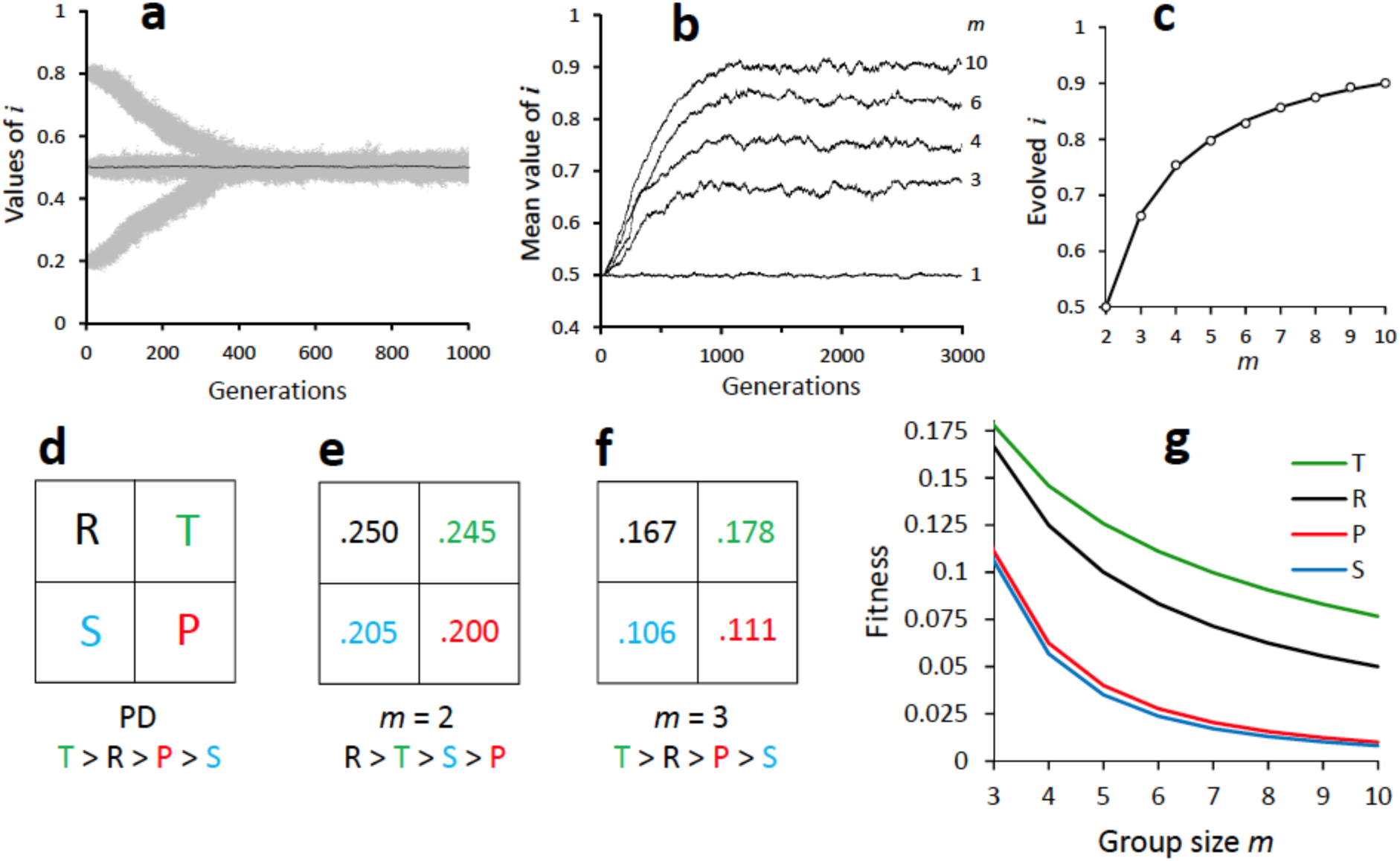
. Linear tradeoff and the evolution of PD. All populations evolved with a linear tradeoff and a Monte Carlo simulation with population size of *N* = 1000, genomic mutation rate of *u* = 0.2 and a Gaussian distribution of mutational effects with mean zero and standard deviation σ = 0.005 (see Materials and Methods for additional details). (**A**) Evolutionary changes with *m* = 1. Grey areas represent all individual *i* values over time in three independent populations started with *i* = 0.8, 0.5 and 0.2. Black trace (—) represents mean *i* values for population started with *i*= ½. A value of *m* = 1 serves as a control for the consequences of clonal selection because all individuals in a group descend from one individual. The consequence of clonal selection is that replication and the production of public goods evolves to the optimum of equaling each other, or *i_k_* = 1 − *i_k_* = ½. (**B**) Traces of mean *i* values for independent populations evolved with increasing values of *m*. (**C**) Match of *i* values predicted by analytical solution *i_a_* = 1 − 1*/m* and mean values evolved by Monte Carlo simulations. (**D**) General payoff matrix representing PD. The payoffs are the fitness values reward *R* when both players cooperate, temptation *T* for one player to defect, sucker’s payoff *S* for the cooperator facing defection, and penalty *P* for both players defecting. PD requires the rank order *T* > *R* > *P* > *S*. (**E**) Fitness payoff matrix for *m* = 2 (see Materials and methods for matrix estimation). The population avoids PD because *R* is the highest value. Cooperation is favored because group size is sufficiently small to allow clonal selection. (**F**) Fitness payoff matrix with *m* = 3. The required PD rank order is satisfied. Optimal cooperation of *i* = ½ is not possible with PD, and defection leads to the evolution of PD and the evolved value of *i_a_* = 1 − 1/*m* (equation (2)). (**G**) Relationship between payoffs *T*, *R*, *P*, and *S* with increasing m. Required rank order for PD is satisfied for all *m* > 2.

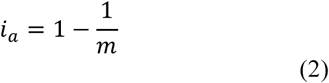
matched closely all of the steady state values evolved with the model (figure 2C). It is also an evolutionarily stable strategy (ESS) because mutants with either higher or lower values of *i* have a lower fitness and cannot invade. Indeed, noticing that *1/m* is equivalent to a coefficient of relatedness between viruses within a host, this result is formally identical to the ESS condition previously derived by Frank [7,8]. Note that if *m* = 2, *i_a_* = *i_k_* = ½ and selection for cooperation is as optimal as kin selection. However, evolution was trapped by PD for *m* ≥3. By constructing the individual fitness of *i_k_* and *i_a_* viruses in pure and mixed groups of size *m* ≥ 3, we found that the resulting fitness values conformed to a PD payoff matrix (figures 2D-F). Thus, our first model with linear tradeoffs remained trapped by the PD domain. Replication effort *i* evolved to high values, while the production of public goods 1 – *i* evolved to low levels. Only clonal or kin selection (*m* = 1) was able to select for high levels of cooperation with 1 – *i* = ½.

### (b) A non-linear fitness tradeoff allows for an evolutionary transition from PD to SD

Our examination of a non-linear tradeoff was motivated by the molecular biology of VSV DI particles described above [13]. Although a linear tradeoff was justified for replication and transcription (figure 1A), DI particles are more than just defectors with *i* = 1 and producing no public goods (1 – *i* = 0). Once a defector evolves to produce zero public goods, it can delete the coding sequences and replace its replication and transcription site with a replication-only site (figure 1B). These changes introduce a non-linear tradeoff because such an ultra-defector is able to make *i* = 1 + *e* copies of its genome, where *e* is the amount gained by having to replicate a smaller genome and not having to spend time on transcription (figure 1C). The value of *e* is a constant representing a cap to the non-linear gain. Thus, *i* is now able to evolve in the interval *i* ∈ [0, 1 + *e*] through mutations. Because it is now also possible that *i* > 1, the new constraint 1 − *i* ≥ 0 was added to the model to prevent public goods from being produced at a negative rate. By incorporating this non-linearity in the fitness tradeoff, we have expanded the scope of models previously proposed by Frank [7,8].

Equation (1) can still be used with the non-linear modifications, although fitness *W_n_* is now determined by two discrete and discontinuous functions. We considered modeling the non-linearity with a single continuous concave up function (*e.g*., exponential or parabolic), but chose to our formulation because we deemed it to be more accurate. When a virus tradeoffs by allocating more effort to either transcription or replication, a linear tradeoff is realistic. However, once tradeoff evolves to allocate *i* = 1 into replication, the virus is free to delete coding regions. As more and more regions are deleted, the replication fitness of the virus increases but its transcription effort remains unchanged because it is already equal to zero. Thus, the fitness tradeoff of a real virus is actually determined by two discrete molecular mechanisms, initially one that affects transcription and later one that does not. A single continuous function would force a tradeoff when there is none, or assume none when there is one.

The addition of a non-linear tradeoff and *i* > 1, altered qualitatively the evolutionary dynamics of our model (figure 3A). As an illustrative example, letting *m* = 8 and *e* = ½, we started with a monomorphic population with *i* = ½ and allowed evolution to proceed with mutations. The population quickly evolved to a steady state of *ia* (equation (2)) as we had reported for a linear model and PD (cf. figure 2B; *m* = 8). However, as mutations accumulated, the population bifurcated into a lineage of ultra-defectors and one of ultra-cooperators. In the ultra-defectors, *i* evolved upwards until it equaled the cap of 1 + *e*, while in the ultra-cooperators it evolved downwards to the clonal selected optimum of *i_k_* = ½ Note that the evolution of ultra-cooperators occurs only after the ultra-defectors begin to increase in frequency. Thus, a mutant ultra-defector must have a minimum value of *i* in order to invade the steady state *i_a_* population. By using equation (1) to estimate the fitness of the DI mutant in the *i_a_* population, and assuming that the mutant is at a low frequency, the minimum *i* valued needed by the ultra-defector was found to be 
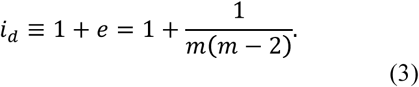
(see section 4c below for the analytical derivation). The bifurcation and the evolution of the ultra-defector also allow the population finally to escape PD and transition into SD. By determining the individual fitness of *i_k_* and *i_d_* viruses in pure and mixed groups of size *m* ≥ 3, there are again only two competitors and the resulting fitness values can be represented by a 2×2 fitness payoff matrix. The results conformed to the SD payoff matrix (figure 3B-C).

**Figure 3.**
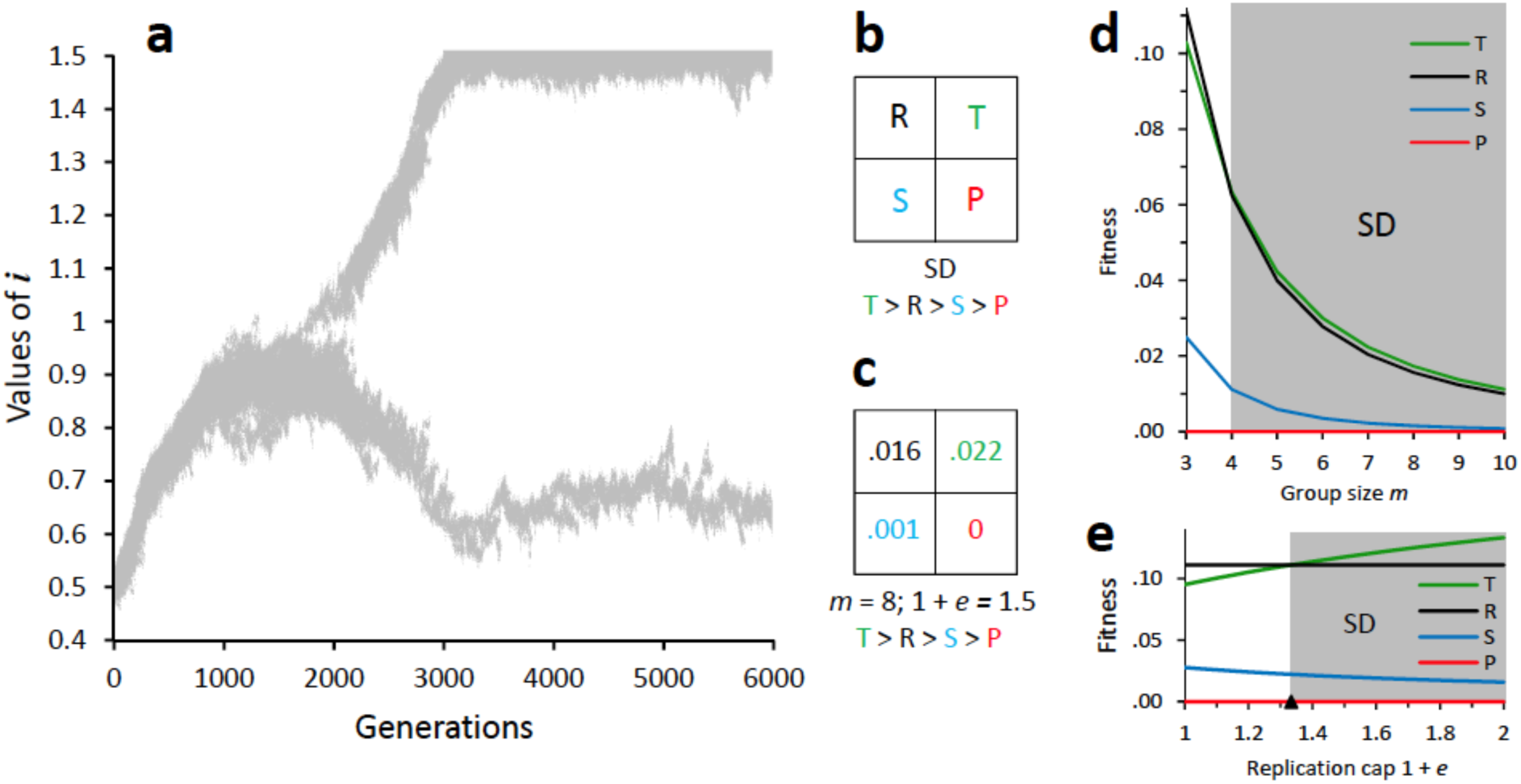
Nonlinear tradeoff and the transition to SD. All populations evolved with a nonlinear tradeoff (*e* > 0) via a Monte Carlo simulation using the same parameters as in figure 2, unless otherwise specified. (**A**) Evolutionary changes with *m* = 8 and a replication cap of 1 + *e* = 1.5. Grey areas represent all individual *i* values over time in a population started with *i* = 0.5. Population evolves steadily higher *i* values to 1 - 1*/m* = 7/8 (equation (2)) and mutations increasing *i* > 1 surface at 1200 generations. At 1800 generations a mutant evolves into an ultra-defector with *i* = 1.5. In response to the evolution of the ultra-defector, the population splits into a second lineage of ultra-cooperators that evolves a lower replication rate *i*, or a higher rate of public good production of 1 − *i*. (**B**) General fitness payoff matrix representing SD with rank *T* > *R* > *S* > *P* (see figure 2 for additional details). (**C**) Payoff matrix for population and conditions in figure 2A (see Materials and methods for matrix estimation). Rank matches requirement for SD. The population is polymorphic and ultra-cooperators and ultra-defectors coexist because ultra-defectors can invade a population of ultra-cooperators (*T* > *R*) and *vice versa* (*S* > *P*). (**D**) Relationship between fitness payoffs *T*, *R*, *S*, and *P* with non-linear tradeoffs 1 + *e* = 1.15 and increasing m. The threshold for evolving SD is given by equation (3) or 1 + *e* > 1 + 1/[*m*(*m* − 2)], which is satisfied for *m* ≥ 4. For *m* = 3, the threshold is not satisfied and the rank is PD because 1.15 < 1.33. (**E**) Relationship between fitness payoffs *T*, *R*, *S*, and *P* with *m* = 3 and increasing replication cap 1 + *e*. With *m* = 3, the threshold for evolving SD is 1.33 (▲) as in figure 2D. The rank is PD for 1 + *e* < 1.33 and SD for 1 + *e* > 1.33.

Equations (2) and (3) can be used to partition parameter space and constructing a landscape for the evolution of cooperation. As the model has only two parameters, let 1 + *e* and *m* be the *y* and *x* axes, in which case equation (3) delineates the boundary between PD and SD (figure 4A). Thus, for all values of *m* ≥ 3, PD evolves if 1 + *e* < *i_d_*, and SD evolves if 1 + *e* > *i_d_*. For *m* values as low as 1 or 2, clonal selection is sufficiently strong and the optimum *i_k_* = ½ evolves. Note that when *e* = 0, the model reverts to the linear form and the outcome is PD as described by equation (2). By plotting the amount of public goods produced as the response variable on the *z* axis, a landscape for the evolved level of cooperation as a function of 1 + *e* and *m* is generated (figure 4B). Public goods are produced maximally by clonal selection, but nearly equivalent amounts are produced by SD. Moreover, as 1 + *e* and *m* are increased in SD, ultra-cooperators are selected to make even more public goods to make up for consumption of the progressively stronger and more numerous ultra-defectors. Indeed, it can be shown mathematically that as 1 + *e* increases, *i_a_* tends to *i_k_* = ½ (see section 4e below). However, because the frequency of individuals producing public goods is not equal (figure 4C), we also examined the landscape for the mean production of public goods as a response variable (figure 4D). Because the frequency is low for high values of 1 + *e* and *m*, the highest mean production, outside of clonal selection regime, is situated centrally, in the SD region, and just beyond the *i_d_* boundary.

**Figure 4.**
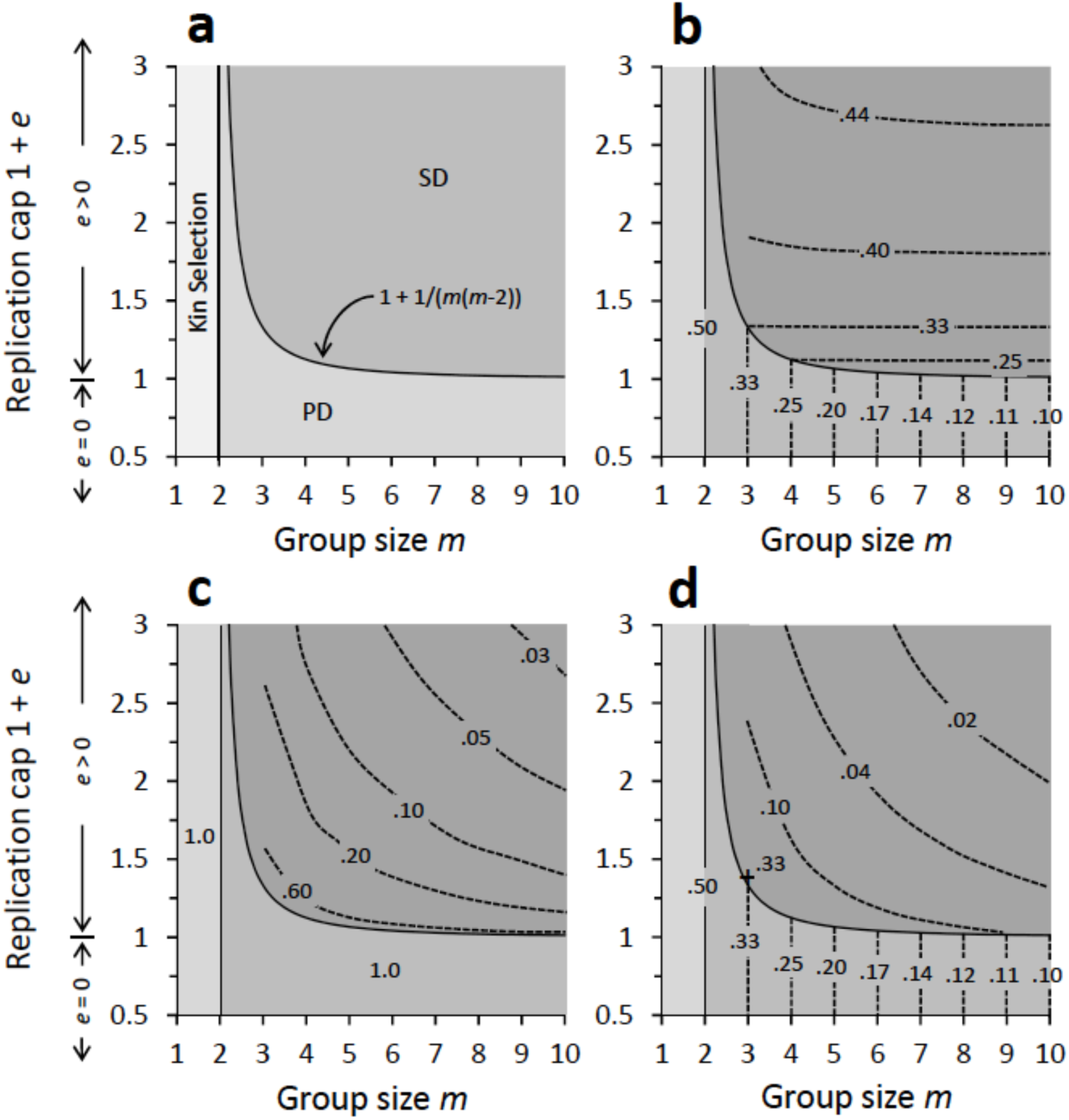
Parameter landscapes for the evolution of cooperation. Plots are topographic representations of evolutionary outcomes projected onto the parameter space of group size *m* and replication cap 1 + *e*. All graphed outcomes were obtained as either analytical or numeric solutions and were also verified through comparisons to Monte Carlo simulations (see Materials and methods). (**A**) Parameter space leading to the evolution of clonal selection, PD, and SD. Boundary for clonal selection and PD is given by equation (2) and for PD and SD by equation (3). (**B**) Evolved maximal level of individual cooperation represented as the production of public goods 1 − *i*. Values in the PD region are from equation (2). Values in SD region are numerical solutions (see Materials and methods) representing the production by ultra-cooperators. (**C**) Frequency of individuals producing the maximal individual values in figure 2B. Clonally-selected and PD populations are monomorphic and all individuals produce maximally. SD populations are polymorphic and frequencies represent ultra-cooperators. (**D**) Evolved mean level of individual cooperation 1 − *i* in populations. Because PD populations are monomorphic and ultra-defectors in SD populations do not produce public goods, the mean is the product of individual values and frequencies from figure 2B-C. A maximum value of 0.33 (+) was observed at *m* = 3 and 1 + *e* = 1.34.

## 3. Concluding remarks

The evolution of cooperation remains a problem partly because it has been easier to identify the barriers to the process rather than solutions. PD presents an obstacle, but it is not clear that escaping PD promotes cooperation. The cooperator could evolve to resist better the defector, but that only makes the goods less public, as did reciprocation and kin selection. SD was recognized as promoting more cooperation [9], but we know little about its evolutionary maintenance, origin, and link to PD and kin or clonal selection. Our model resolves many of these issues by mapping the evolution of cooperation onto two parameters, group size and the magnitude of the non-linear gain to an ultra-defector. On this parametric space, increasing group size alone is sufficient for driving the evolution of cooperation from clonal (kin) selection to PD and to SD. We show that the transition to SD results from the splitting of the population into lineages of ultra-defectors and ultra-cooperators. However, the split is triggered by the initial evolution of more defection, rather than more cooperation. If the tradeoff between defection and cooperation is linear, PD traps the population because the evolution of more defection, and the split, is prevented. With a non-linear tradeoff, more defection and the ultra-defector are able to evolve. Once the ultra-defector increases in frequency, the cooperator is selected to make even more public goods and to evolve into the ultra-cooperator of SD. The best ultra-cooperators make nearly as much public goods as clonally-selected individuals. The numerical analysis of equation (10) supports this latter conclusion: for any given value of *m* one can explore which values should take *i* to hold the equilibrium frequency of wildtypes, 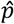, constant if the DIs take increasingly large values of 1 + *e*. It turns out that at the limit when 1 + *e* →∞, then *i* → ½ This means that the optimal solution for a wildtype virus in presence of an ultra-defector DI would be to become itself and ultra-cooperator and invest 50% of its resources in the production of common goods.

The effect of increasing group size on the evolution of cooperation has been observed in laboratory studies with RNA viruses. The first transition from clonal selection to PD was seen in the bacteriophage φ6, where cheaters spontaneously arose when group size was increased to *m* = 5 [10,11,32]. This was a reversible process, as cheaters were outcompeted when a regime of clonal selection (*m* << 1) was imposed [11]. The second transition, from PD to SD, has been documented in many RNA viruses by the evolution of DIs in dense cultures when large viral groups are able to co-infect the same host cell [33-37]. Subsequent evolution in some of these dense cultures often result in population cycles during which wildtype viruses become resistant to the effects of the DIs [18,19,38-41] and the DIs evolve in turn the ability to overcome the resistance [15,16,39-41]. Such cycles are driven by the evolution of new wildtype RNA replicases that no longer recognize the replication signals of the DIs, followed by the co-evolution of DIs with altered signals that are recognized [39]. Because the aim of our model was to analyze the evolutionary dynamics of cooperation between a given pair of wildtype and DI, such co-evolutionary races for novel replicases were not considered. In other words, our model focuses on the period of stasis between cycles. Periods of stasis can be long, and evolutionarily important, because the start of a new cycle requires a double mutation in the wildtype virus. One mutation must change the replicase to provide resistance, but a second one is needed to change the wildtype replication signal so that it recognizes the mutated replicase. Double mutants will be rare and less likely to appear if population size is small. While laboratory populations can be large, wild populations may be much smaller because of bottlenecks or other ecological stresses.

Although our model was motivated by viral biology, there are clear overlaps between our approach and previous studies. At the mathematical and the theoretical level, viruses, prebiotic replicons, and humans can sometimes be modeled equivalently [4,5,7-9]. SD was recognized as favoring the evolution of higher cooperation levels [8], but because the games had been studied as separate games played in isolation, there was no evolutionary connection between them. For a population to evolve more cooperation by transitioning from PD to SD, the payoff matrix had to be changed. However, the rules controlling the matrices and their changes were not apparent, or at least assumed to be too complex to be derived from known ecological and evolutionary processes. Our finding that varying group size alone, so long as the tradeoffs between replication and the production of public goods is non-linear, controls the transition between PD and SD, and greatly simplifies the conditions required for the evolution of cooperation. Our payoff matrices were able to evolve freely, and the ones emerging as ESS conformed to PD and SD as group size changed. Because all organisms experience a group size, the results of our model show that the evolution of cooperation may not need much more than changing one ecological parameter. We hope that our work will stimulate additional work on RNA viruses or other organisms that produce public goods as model systems to explore social evolution.

## 4. Materials and methods

### (a) Monte Carlo model for simulating of evolution in viral populations

A population of size *N* viruses was constructed by assigning a starting value of *i* to each virus. The assigned value could be randomly or deliberately chosen to explore different starting scenarios. The population was then divided into *N/m* groups of size *m* and the fitness of individual viruses was determined with equation (1). To create the population for the next generation, the current population was sampled *N* times with replacement to ensure a constant population size. The sampling was biased by using the normalized fitness values of each virus as their probability of being chosen. Fitness was determined using equation (1) with the appropriate linear and non-linear values of *i* and 1 − *i*. This sampling bias introduces natural selection and evolution could thus be modeled and followed over generations. Whenever desired, mutations were introduced by changing individual values of *i* with a probability of 0 < *u* < 1. If mutations were not desired, a value of *u* = 0 was used instead. If a virus was to be mutated, its *i* value was changed by an amount randomly drawn from a Gaussian distribution with a mean zero and a specified standard deviation, σ. All simulations were coded in R version 3.2.4 computer language.

### (b) Analytical solution of equation (2)

Let *j* and *i* be the replication effort of a mutant and wildtype virus, and *j* > i. The fitness of a wildtype virus in a population all formed by wildtype viruses can be computed using equation (1). In such case the burst size would be 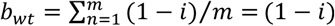 and the total replication effort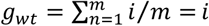 because the coinfection group has a uniform composition. Thus,

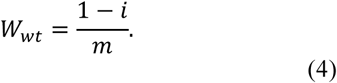

Now let’s imagine that a defector mutant that invests *j* > *i* into replication appears in a population of *m* – 1 wildtype viruses. The fitness of this mutant can be calculated using equation (1) but now considering that *b_def_* = [(*m* − 1)(1 − *j*)]*/m* and *g_def_* = [(*m* − 1)*i*+*j*]*m*: 
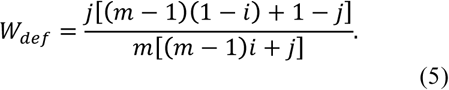

Coexistence of both genotypes would occur if, and only if, *W_wt_* = *W_def_*. If *W_wt_* < *W_def_*, the defector mutant will invade, whereas the opposite condition means that invasion is not possible. Combining equations (4) and (5) and simplifying, we found that there are two different values of *j* that satisfy the coexistence condition: *j* = *i* and *j* = (1 − *i*)(*m* − 1). These two lines correspond to pairs of values (*i, j*) in which defector mutant and wildtype have equal fitness. The intersection of these two fitness isoclines results in the equilibrium condition shown in equation (2): 
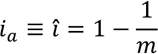

The linear stability of this fixed point was evaluated by constructing the Jacobian matrix of the system formed by equations (4) and (5), which is given by 
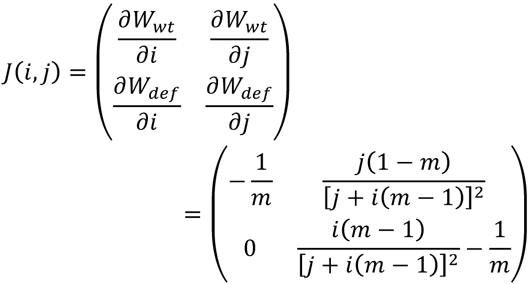

The two eigenvalues for the fix point (*î*,*ĵ*) are *λ*_1_ + = *1/m*^3^ and *λ*_2_ = − 1/*m*. Since *λ*_1_ > *0* and *λ_2_* < *0* ∀ *m* > 0, the fix point takes the form of a saddle and is thus instable.

### (c) Analytical solution of equation (3)

The solution is obtained by exploring the conditions in which a DI mutant will invade a population at the equilibrium specified by equation (2). Substituting *i* by the equilibrium condition given by equation (2) into equations (4) and (5) and recalling that DIs do not contribute to the production of common goods and have an investment in reproductive effort 1 + *e* > *i*, we obtain after some algebraic work the following fitness equations for wildtype and DI, respectively: 
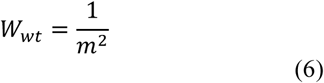
and 
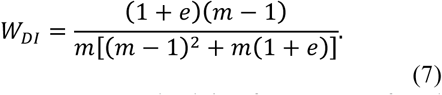

By making *W_wt_* = *W_DI_* and solving for 1 + *e*, we found the 1 + *e* value at which wildtype and DI viruses will coexist (equation (3)): 
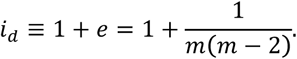

### (d) Constructing payoff matrixes for PD and SD

Payoff values in a game theory matrix traditionally represent interactions between two individuals rather than between many in a large group. Because our models consider groups of size *m* that can be much larger than two, we adapted the payoff matrix 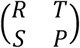 to represent individual fitness values in populations with only cooperators, only defectors, one cooperator invading a large population of defectors, and one defector invading a large population of cooperators. The fitness of the invading cooperator, which equals the value the virus has when it is alone in a group with *m* – 1 defectors, represents *S*. The fitness of the invading defector represents *T*, or the value when it is alone in a group of *m* – 1 cooperators. The fitness of a cooperator in a population with only cooperators represents *R*, or the value when it is in a group with *m* cooperators. The fitness of a defector in a population with only defectors represents *P*, or the value when it is in a group with *m* defectors. A payoff matrix adapted to larger groups retains the predictive properties for an ESS analysis. If *T* > R, a defector is able to invade a population of cooperators. If *S* > *P*, a cooperator is able to invade a population of defectors, which is one of the requirements for SD. All fitness values were based on equation (1) and using the appropriate linear and non-linear values of *i* and 1 − *i*.

Although we can have potentially have *m* players in a group, the interaction can be reduced to a two-player game for the construction of a 2×2 playoff matrix. In an ESS analysis, the resident population (even if it is polymorphic) is always held constant, in which case the two players are the resident(s) and the invader.

### (e) Equilibrium frequencies of DI and wildtype viruses in SD

With SD, the tradeoff is necessarily non-linear and both DIs and wildtype viruses are present in the population. We calculate here the equilibrium frequencies of DI and wildtype viruses in the population for a given set of values for group size *m*, DI replication fitness 1 + *e*, and wildtype replication fitness *i*.

Let *p* and 1 – *p* represent, respectively, the frequency of wildtype and DI viruses in the population. Assuming that the viruses are distributed by a binomial process into groups of size *m*, the probability of getting x wildtypes is *P*(*x*)=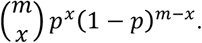. The mean individual fitness of the viruses in the population is then given by

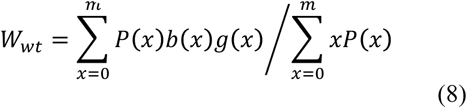
and
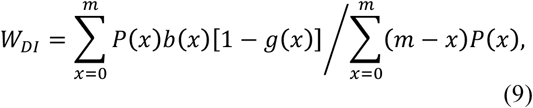
where *b*(*x*) = *x*(*1* − *i*)*/m* is the total production of common goods by a group of size *m* that contains *x* wildtypes, and *g*(*x*) = *xi/*[*xi* + (*m* − *x*)(1 + *e*)] is the relative individual replication effort by the wildtype virus in the same group. Because DIs do not make public goods, they do not contribute but have access to *b*(*x*). On the other hand, because DIs replicate, their relative individual replication rate is 1 – *g*(*x*).

Noting that the denominators of equations (8) and (9) are the expectations of the binomial distribution *mp* and *m*(1 – *p*), the frequency of DIs and wildtypes will be unchanging and at their equilibria 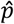 and 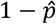 when *W_wt_* = *W_DI_*, or 
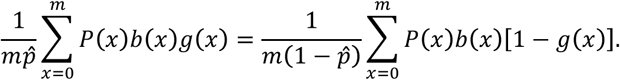

After replacing *P*(*x*), *b*(*x*) and *g*(*x*) by their actual values, we obtain the following expression:

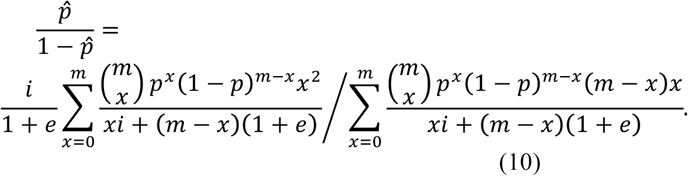

Equation (10) can be solved analytically for any value of *m*≥2 to obtain the equilibrium frequency of the cooperator virus as a function of *i* and *e*, namely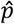(*i*,*e*|*m*). For instance, in the case *m* = 3, the equilibrium frequency for the cooperator virus is given by 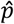 (*i*,*e*|3) = *i*(1 + *e* + 2*i*)/[2(1 + *e*)(1 + e – *i*)], which takes positive values for any *i* < 1 + *e* value. Linear stability analysis shows that the two eigenvalues of the Jacobian matrix *J*(*i*,*e*) = 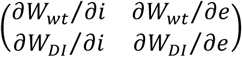 evaluated at the fix point 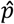 (*i*,*e*|3) are + *λ*_1_ < 0 and *λ*_2_ > *0*, as corresponds to a unstable saddle fix point. For *m* ≥ 4, *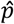*(*i*,*e\m*) is a complex polynomic function of *i* and *e*, with at least one root in the interval [0, 1] that in every case corresponds to an unstable saddle point. Equation (10) can also be solved numerically for every value of *m* to find the frequency *p* of wildtypes in the population as a function of values of *i* and *e*.

### (f) Numerical solutions for the evolved values of replication fitness *i* and of the frequency of ultra-cooperators in an SD parameter space

From equation (10), we first obtained the equilibrium frequency of wildtype and DI viruses in a population, 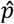 and 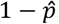, for a given group size of *m*, a DI replication fitness of 1 + *e*, and a wildtype replication fitness of *i*. These equilibrium values are not necessarily the ones that will evolve by natural selection in a population. To determine if they could be the evolved values, we evaluated whether a population with these frequencies of 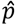 and 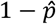 could be invaded by a mutant wildtype virus. Invasion by mutant DI was not examined because we had found in our Monte Carlo simulations that 1 + *e* would evolve to be infinitely large and thus had to be capped at its chosen value (figure 3a). The capping is justified on the basis of physiological limit to replication speed in the host cell. The evolution of the wildtype replication fitness was frequency dependent, which is why we need to evaluate by an invasion criterion and searching for an ESS value.

To assess the invasion, we first estimated the individual fitness of DI and wildtype viruses distributed by a binomial process in groups of size *m*. By knowing the binomial composition of each group, the individual fitness of DI and wildtype viruses in the group could be determined with equation (1). To test invasiveness, we then introduced one mutant wildtype with a replication fitness of *i* ± *δ*. The individual fitness of the mutant was estimated by adding it to all binomially distributed groups of size *m* – 1. The mutant was judged to be able to invade if it had an individual fitness, again estimated by equation (1), greater than the wildtype virus. A value of *δ* = 0.001 was used for all evaluations. The evolved value of *i* was found by searching over a range of [0, 1] a value that was not invasible. The 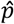 and 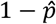 values corresponding to the evolved *i* were then used as the evolved frequencies of the DI and wildtype viruses. These evolved values are depicted in the SD parameter space of figures 4B-D. Estimates of these evolved values were also obtained from our Monte Carlo simulations. Our numerical and Monte Carlo estimates were highly correlated (see figure S1, Supplementary material). The search code was written in R version 3.2.4 computer language.

## Acknowledgements

We dedicate this paper to the memory of John J. Holland, who first brought the authors to UCSD and taught us so much about defective viral genomes and virus evolution.

## Author contributions

LC and SFE, Conception and design, Acquisition of data, Analysis and interpretation of data, Drafting or revising the article.

## Supplementary Material

**Figure S1.**
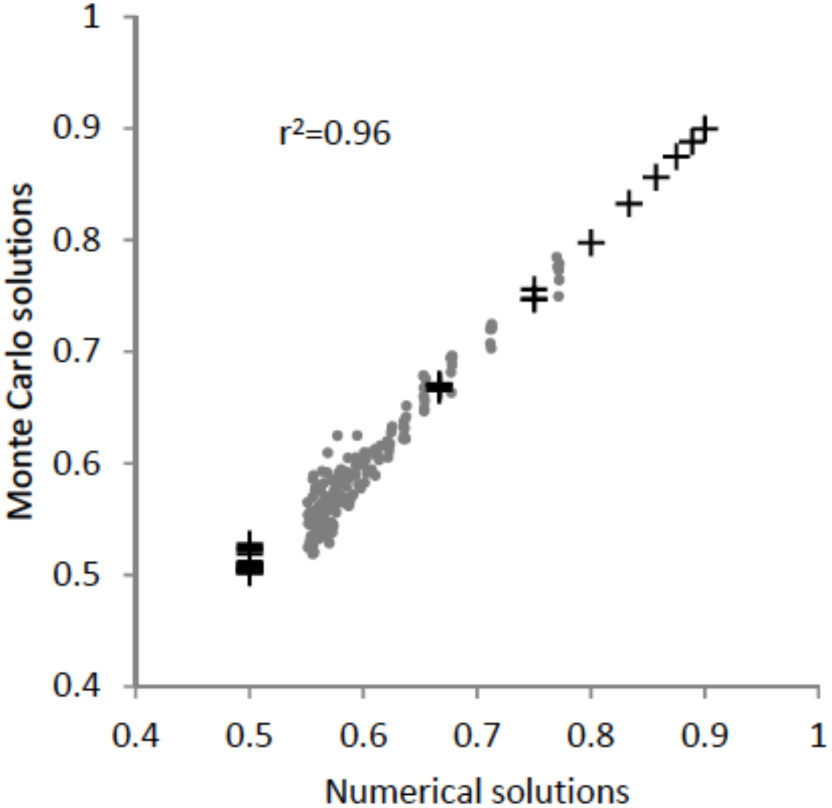
Comparing Monte Carlo and numerical solutions for evolved values of *i*. Solutions obtained as described in Materials and Methods. Individual points represent *i* values obtained by the Monte Carlo (MC) and numerical solutions. MC simulations were run for 10,000 generations with population size of *N* = 500*m*, where *m* is group size, and a mutation rate of *u* = 0.2 and a Gaussian distribution of mutational effects with mean zero and standard deviation σ = 0.005. Populations generally reach equilibrium values after 1000 generations (figure 2A). Reported values of *i* are the mean value at the last generation. The parameter space represented in figures 4A-D was explored by letting *m* range from 2, 3, 4,…,10 and 1 + *e* from 1, 1.1, 1.2,…, 3.0. Values of *i* corresponding to PD (+) and SD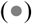. The match between MC and numerical solutions yielded a value of *r*^2^ = 0.96.

## References

1. Bergua M, Zwart MP, El-Mohtar C, Shilts T, Elena SF, Folimonova SY. 2014 A viral protein mediates superinfection exclusion at the whole organism level but is not required for exclusion at the cellular level. J. Virol. 88, 11327–11338. (doi: 10.1128/JVI.01612-14)

2. Turner PE, Burch CL, Hanley KA, Chao L. 1999 Hybrid frequencies confirm limit to coinfection in the RNA bacteriophage φ6. J. Virol. 73, 2420–2424.

3. Whitaker-Dowling PA, Youngner JS, Widnell CC, Wilcox DK. 1983 Superinfection exclusion by vesicular stomatitis virus. Virology 131, 137–143.

4. Doebeli M, Hauert C, Killingback T. 2004 The evolutionary origin of cooperators and defectors. Science 306, 859–862. (doi: 10.1126/science.1101456)

5. Doebeli M, Hauert C. 2005 Models of cooperation based on the Prisoner's Dilemma and the Snowdrift game. Ecology Lett. 8, 748–766. (doi: 10.1111/j.1461-0248.2005.00773.x)

6. Elena SF, Bernet GP, Carrasco JL. 2014 The games plant viruses play. Curr. Opin. Virol. 8, 62–87. (doi: 10.1016/j.coviro.2014.07.003)

7. Frank SA. 1994 Kin selection and virulence in the evolution of protocells and parasites. Proc. R. Soc. B 258, 153–161. (doi: 10.1098/rspb.1994.0156)

8. Frank SA. 1995 Mutual policing and repression of competition in the evolution of cooperative groups. Nature 377, 520–522. (doi: 10.1038/377520a0)

9. Kummerli R, Colliard C, Fiechter N, Petitpierre B, Russier F, Keller L. 2007 Human cooperation in social dilemmas: comparing the Snowdrift game with the Prisoner's Dilemma. Proc. R. Soc. B 274, 2965–2970. (doi: 10.1098/rspb.2007.0793)

10. Turner PE, Chao L. 1999 Prisoner's dilemma in an RNA virus. Nature 398, 441–443. (doi: 10.1038/18913)

11. Turner PE, Chao L. 2003 Escape from Prisoner’s Dilemma in RNA phage φ6. Am. Nat. 161, 497–505. (doi: 10.1086/367880)

12. Dimmock NJ. 1991 The biological significance of defective interfering viruses. Rev. Med. Virol. 1, 165–176.

13. Holland JJ. 1986 Generation and replication of defective viral genomes. In Fields BN, Knipe DM, editors-Fundamental Virology. New York: Raven Press. p 6.77–6.99.

14. Szathmáry, E. 1992 Natural selection and dynamical coexistence of defective and complementing virus segments. J. Theor. Biol. 157, 383–406. (doi: 10.1016/S0022-5193(05)80617-4)

15. Kirkwood TBL, Bangham CRM. 1994 Cycles, chaos, and evolution in virus cultures: a model of defective interfering particles. Proc. Natl. Acad. Sci. USA 91, 8685–8689. (doi: 10.1073/pnas.91.18.8685)

16. Bangham CRM, Kirkwood TBL. 1990 Defective interfering particles: effects in modulating virus growth and persistence. Virology 179, 821–826. (doi: 10.1016/0042-6822(90)90150-P)

17. Cole CN, Baltimore D. 1973 Defective interfering particles of poliovirus. 3. Interference and enrichment. J. Mol. Biol. 76, 345–361. (doi: 10.1016/0022-2836(73)90509-3)

18. Giachetti C, Holland JJ. 1988 Altered replicase specificity is responsible for resistance to defective interfering particle interference of an Sdi-mutant of vesicular stomatitis virus. J. Virol. 62, 3614–3621.

19. Giachetti C, Holland JJ. 1989 Vesicular stomatitis virus and its defective interfering particles exhibit in vitro transcriptional and replicative competition for purified L-NS polymerase molecules. Virology 170, 264–267. (doi: 10.1016/0042-6822(89)90375-9)

20. Horodyski FM, Holland JJ. 1984 Reconstruction experiments demonstrating selective effects of defective interfering particles on mixed populations of vesicular stomatitis virus. J. Gen. Virol. 65, 819–823. (doi: 10.1099/0022-1317-65-4-819)

21. Kolakofsky D. 1976 Isolation and characterization of Sendai virus DI-RNAs. Cell 8, 547–555. (doi: 10.1016/0092-8674(76)90223-3)

22. Nayak DP, Sivasubramanian N, Davis AR, Cortini R, Sung J. 1982 Complete sequence analyses show that two defective interfering influenza viral RNAs contain a single internal deletion of a polymerase gene. Proc. Natl. Acad. Sci. USA 79, 2216–2220. (doi: 10.1073/pnas.79.7.2216)

23. Nonoyama M, Graham AF. 1970 Appearance of defective virions in clones of reovirus. J. Virol. 6, 693–694.

24. Stampfer M, Baltimore D, Huang AS. 1969 Ribonucleic acid synthesis of vesicular stomatitis virus. I. Species of ribonucleic acid found in Chinese hamster ovary cells infected with plaque-forming and defective particles. J. Virol. 4, 154–161.

25. Weiss B, Goran D, Cancedda R, Schlesinger S. 1974 Defective interfering passages of Sindbis virus: nature of the intracellular defective viral RNA. J. Virol. 14, 1189–1198.

26. López CB. 2014 Defective viral genomes: critical danger signals of viral infections. J. Virol. 88, 8720–8723. (doi: 10.1128/JVI.00707-14)

27. Frensing T. 2015 Defective interfering viruses and their impact on vaccines and viral vectors. Biotechnol. J. 10, 681–689. (doi: 10.1002/biot.201400429)

28. Notton T, Sardanyés J, Weinberger AD, Weinberger LS. 2014 The case for transmissible antivirals to control population-wide infectious diseases. Trends Biotechnol. 32, 400–405. (doi:10.1016/j.tibtech. 2014.06.006)

29. Dixit NM, Perelson AS. 2004 Multiplicity of human immunodeficiency virus infections in lymphoid tissue. J. Virol. 78, 8942–8945. (doi: 10.1128/JVI.78.16.8942-8945.2004)

30. Gutiérrez S, Yvon M, Thébaud G, Monsion B, Michalakis Y, Blanc S. 2010 Dynamics of the multiplicity of cellular infection in a plant virus. PLoS Pathog. 6, e1001113. (doi: 10.1371/jounal.ppat.1001113)

31. Tromas N, Zwart MP, Lafforgue G, Elena SF. 2014 Within-host spatiotemporal dynamics of plant virus infection at the cellular level. PLoS Genet. 10,e1004186. (doi: 10.1371/pgen. 1004186)

32. Turner PE, Chao L. 1998 Sex and the evolution of intrahost competition in RNA virus φ6. Genetics 150, 523–532.

33. Akpinar F, Inankur B, Yin, J. 2016 Spatial-temporal patterns of viral amplification and interference initiated by a single infected cell. J. Virol. 90, 7552–7566. (doi: 10.1128/JVI.00807-16)

34. Bangham CRM, Kirkwood TBL. 1993 Defective interfering particles and virus evolution. Trends Microbiol. 1, 260–264.

35. Jacobson S, Dutko FJ, Pfau CJ. 1979 Determinants of spontaneous-recovery and persistence in MDCK cells infected with lymphocytic choriomeningitis virus. J. Gen. Virol. 44, 113–121. (doi: 10.1099/0022-1317-44-1-113)

36. Stauffer Thompson KA, Yin J. 2010 Population dynamics of an RNA virus and its defective interfering particles in passage cultures. Virol. J. 7, 257. (doi: 10.1186/1743-422X-7-257)

37. Timm C, Akpinar F, Yin J. 2014 Quantitative characterization of defective virus emergence by deep sequencing. J. Virol. 88, 2623–2632. (doi: 10.1128/JVI.02675-13)

38. Brinton MA, Fernandez AV. 1983 A replication-efficient mutant of West Nile virus is insensitive to DI particle interference. Virology 129, 107–115. (doi: 10.1016/0042-6822(83)90399-9)

39. DePolo NJ, Giachetti C, Holland JJ. 1987 Continuing coevolution of virus and defective interfering particles and of viral genome sequences during undiluted passages–virus mutants exhibiting nearly complete resistance to formerly dominant defective interfering particles. J. Virol. 61, 454–464.

40. Kawai A, Matsumoto S. 1977 Interfering and noninterfering defective particles generated by a rabies small plaque variant virus. Virology 76, 60–71. (doi: 10.1016/0042-6822(77)90282-3)

41. Zwart MP, Pijlman GP, Sardanyés J, Duarte J, Januário C, Elena SF. 2013 Complex dynamics of defective interfering baculoviruses during serial passage in insect cells. J. Biol. Phys. 39, 327–342. (doi: 10.1007/s10867-013-9317-9)

